## supplemental file for "*Wuschel2* enables highly efficient CRISPR/Cas-targeted genome editing during rapid *de novo* shoot regeneration in sorghum"

### Supplementary information

**a** pPHP79066

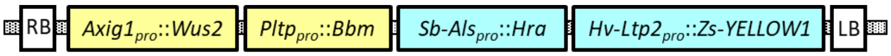


**b** pPHP8181

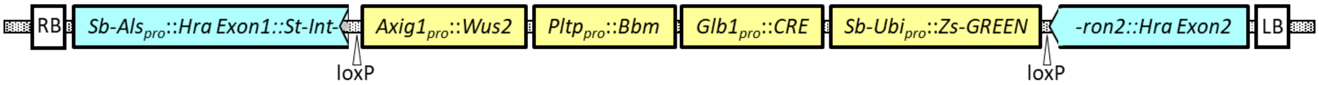


**c** pPHP94632

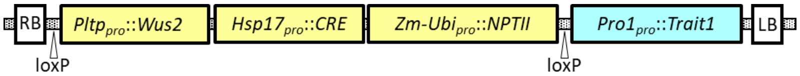


**d** pPHP94292

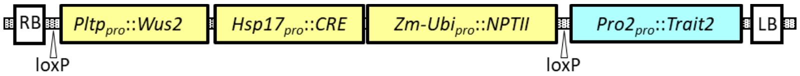


**e** PHP87078 (“Altruistic”)

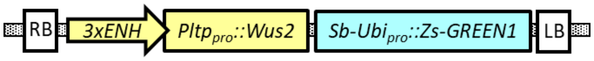


**f** pPHP88158 (“Altruistic”)

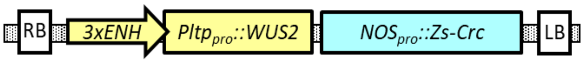


**g** pPHP96564

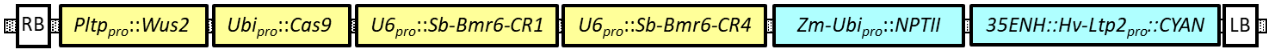


**h** pPHP86655

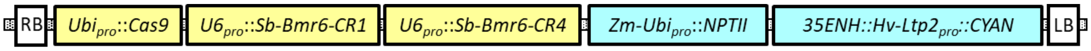


**i** pPHP87980

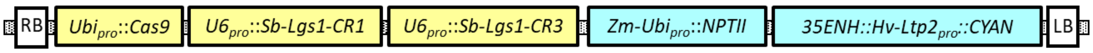

**j** pPHP86801

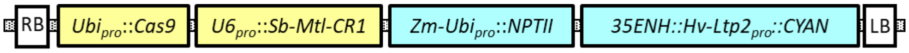


**k** pPHP87098

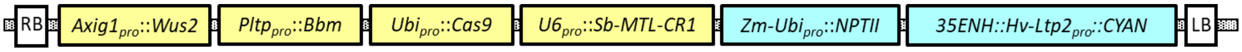


**l** pPHP87018

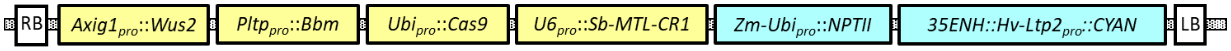


**m** pPHP87984

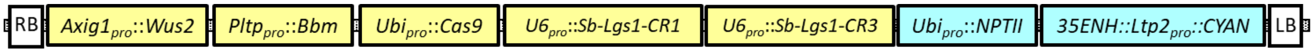


**Supplementary Figure 1| Schematic representation of the molecular components of constructs used in this study**.

### Transformation procedures

**Infection:** Inoculate immature embryos with *Agrobacterium* LBA4404 Thy-harboring ternary vector system in infection medium (**Supplementary Table 1**)

↓

**Co-cultivation:** Cultivate immature embryos with *Agrobacterium* on co-cultivation medium (**Supplementary Table 2**) at 25°C overnight followed by 28°C for 6 days in darkness

**Conventional**

**Callus proliferation w/o selection:** Induce embryogenic callus on the multi-purpose medium (**Supplementary Table 3**) at 28°C in darkness for a week

↓

**Callus proliferation w/ selection:** Continue callus development on callus induction medium (**Supplementary Table 4**) with proper selection reagent at 28°C in darkness for total 9 weeks with subculturing every 3 weeks.

**Morphogenic gene-enabled**

**Somatic embryo formation w/o selection:** Induce somatic embryo formation on the multi-purpose medium (**Supplementary Table 3**) at 28°C in darkness for a week

***Wus2/CRE*-enabled marker-free**

**Somatic embryo formation w/ selection:** Induce somatic embryo formation on the multi-purpose medium (**Supplementary Table 3**) with 250 mg/l G418 at 28°C in darkness for 3 weeks

↓

**Heat-shock:** Incubate at 45°C for 3h

↓

**Maturation w/o selection:** Shoot regeneration on selection-free maturation medium (**Supplementary Table 5**) at 28°C in darkness for 4 weeks

**Maturation w/ selection:** Shoot regeneration on maturation medium (**Supplementary Table 5**) with proper selection reagent at 28°C in darkness for 4 weeks

↓

**Rooting:** Root regeneration on selection-free rooting medium (**Supplementary Table 6**) under light for 1-3 weeks

↓

Rooted plants transferred to greenhouse

**Supplementary Figure 2| Flow diagram of the *Agrobacterium*-mediated transformation systems.**

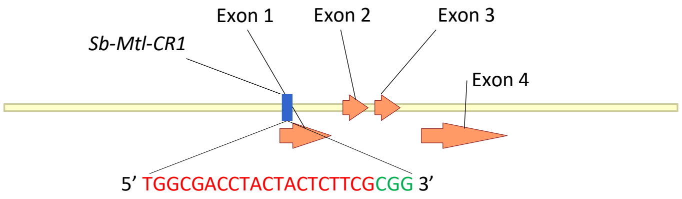


**Supplementary Figure 3| Diagram of the *Sb-Mtl* gene structure and target site.** Gene structure of *Sb-Mtl* with sgRNA target site (in red) and PAM sequence (in green).

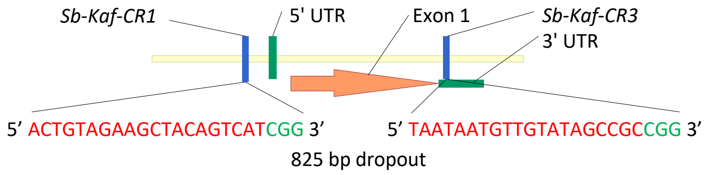


**Supplementary Figure 4| Diagram of the *Sb-Kaf* gene structure and target sites.** Gene structure of *Sb-Kaf* with sgRNA target sites (in red) and PAM sequences (in green).

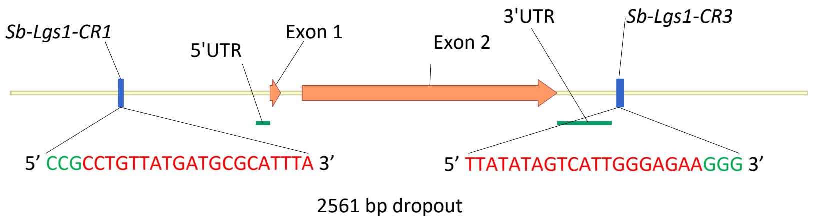


**Supplementary Figure 5| Diagram of the *Sb-Lgs1* gene structure and target sites.** Gene structure of *Sb-Lgs1* with sgRNA target sites (in red) and PAM sequences (in green).

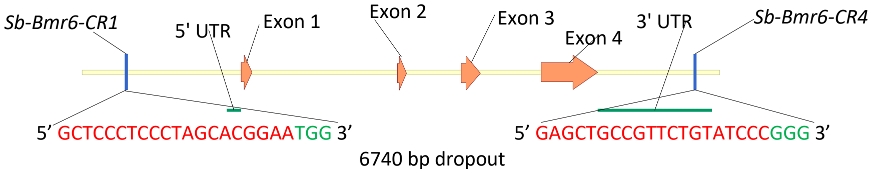


**Supplementary Figure 6| Diagram of the *Sb-Bmr6* gene structure and target sites.** Gene structure of *Sb-Bmr6* with sgRNA target sites (in red) and PAM sequences (in green).

### Supplementary Table 1| Infection medium

| | |
|---|---|
| MS salt | 4.3 g/l |
| Myo-Inostitol | 0.1 g/l |
| Nicotinic acid | 0.5 mg/l |
| Pyridoxine HCl | 0.5 mg/l |
| Thiamine HCl | 1 mg/l |
| Casamino acids | 1 g/l |
| Dichlorophenoxyacetic acid (2,4-D) | 1.5 mg/l |
| Sucrose | 68.5 g/l |
| Glucose | 36 g/l |
| Thymidine | 50 mg/l |
| Acetosyringone | 100 µM (Add fresh) |

### Supplementary Table 2| Co-cultivation medium

| | |
|---|---|
| MS salt | 4.3 g/l |
| Myo-Inostitol | 0.1 g/l |
| Nicotinic acid | 0.5 mg/l |
| Pyridoxine HCl | 0.5 mg/l |
| Thiamine HCl | 1 mg/l |
| Dichlorophenoxyacetic acid (2,4-D) | 2 mg/l |
| Sucrose | 20 g/l |
| Glucose | 10 g/l |
| L-Proline | 0.7 g/l |
| MES Buffer | 0.5 g/l |
| Sigma Agar | 8 g/l |
| Acetosyringone | 100 µM (Add fresh) |
| Ascorbic acid | 10 mg/l |
| Thymidine | 50 mg/l |

### Supplementary Table 3| Multi-purpose medium

| | |
|---|---|
| MS salt | 4.3 g/l |
| Myo-Inostitol | 0.25 g/l |
| Casein hydrolysate | 1 g/l |
| Thiamine HCl | 1 mg/l |
| Dichlorophenoxyacetic acid (2,4-D) | 2 mg/l |
| Maltose | 30 g/l |
| L-Proline | 0.69 g/l |
| Cupric sulfate | 1.22 mg/l |
| Phytagel | 3.5 g/l |
| BAP | 0.5 mg/l |
| Carbenicillin | 100 mg/l |
| Selection reagent* | |

*Either no selection or 250mg/l G418 or 0.1mg/l imazapyr.

### Supplementary Table 4| Callus proliferation medium

| | |
|---|---|
| MS salt | 4.3 g/l |
| Myo-Inostitol | 0.25 g/l |
| Casein hydrolysate | 1 g/l |
| Thiamine HCl | 1 mg/l |
| Dichlorophenoxyacetic acid (2,4-D) | 1 mg/l |
| Maltose | 30 g/l |
| L-Proline | 0.69 g/l |
| Cupric sulfate | 1.22 mg/l |
| Phytagel | 3.5 g/l |
| BAP | 0.5 mg/l |
| Carbenicillin | 100 mg/l |
| Selection reagent* | |

* Either no selection or 250mg/l G418 or 0.1mg/l imazapyr.

### Supplementary Table 5| Maturation medium

| | |
|---|---|
| MS salt | 4.3 g/l |
| Myo-Inostitol | 0.1g/l |
| MS vitamins stock (0.1 g/l nicotinic acid, 0.1 g/l pyridoxine HCl, 0.02 g/l thiamine HCl, 0.4 g/l glycine) | 5 ml/l |
| zeatin | 0.5 mg/l |
| Cupric sulfate | 1.25 mg/l |
| L-Proline | 0.7 g/l |
| Sucrose | 60 g/l |
| Phytagel | 3.5 g/l |
| Indole-3-acetic acid | 1 mg/l |
| ABA | 0.1 µM |
| Thidiazuron | 0.1 mg/l |
| Carbenicillin | 100 mg/l |
| Selection reagent* | |

* Either no selection or 250mg/l G418 or 0.1mg/l imazapyr.

### Supplementary Table 6| Rooting medium

| | |
|---|---|
| MS salt | 2.15 g/l |
| Myo-Inostitol | 0.05 g/l |
| MS vitamins stock (0.1 g/l nicotinic acid, 0.1 g/l pyridoxine HCl, 0.02 g/l thiamine HCl, 0.4 g/l glycine) | 2.5 ml |
| Sucrose | 20 g/l |
| Phytagel | 3 g/l |

### Supplementary Table 7| The integrated morphogenic gene-enabled transformation efficiencies of six Corteva elite lines

| Genotype | # of embryos infected | # of T0 plants | Transformation efficiency (%) |
|---|---|---|---|
| PH1679 | 280 | 4 | 1.4 |
| PH1719 | 233 | 47 | 20.1 |
| PH1181 | 240 | 16 | 6.7 |
| PH2517 | 162 | 29 | 17.9 |
| PH1481 | 235 | 54 | 23 |
| PH2545 | 226 | 56 | 24.7 |

### Supplementary Table 8| *Sb-Mtl* mutation frequency using single gRNA in Tx430

| Transformation system | Construct | sgRNA | # T0 plants analyzed | # of T0 events w/ mutation (freq.) |
|---|---|---|---|---|
| Conventional | pPHP86801 | *Sb-Mtl-CR1* | 29 | 8 (27.5%) |
| Integrated morphogenic gene-enabled | pPHP87098 (*Axig1*$_{pro}$*:Wus2/Pltp*$_{pro}$*:Bbm*) | *Sb-Mtl-CR1* | 97 | 72 (74.2%) |

### Supplementary Table 9| *Sb-Kaf* mutation and gene-dropout frequencies using two gRNAs in Tx430

| Transformation system | Construct | sgRNA | # T0 plants analyzed | # of T0 events w/ mutation (freq.) | # T0 events w/ gene-dropout (freq.) |
|---|---|---|---|---|---|
| Conventional | pPHP87018 | *Sb-Kaf-CR1* | 33 | 13 (39.3%) | 0 |
| | | *Sb-Kaf-CR3* | | 16 (48.5%) | |
| Non-integrated *Wus2*-enabled altruistic | (90%) pPHP87018+(10%) PHP88158 (*3xENH:Pltp*$_{pro}$*:Wus2/NOS*$_{pro}$*:Crc*) | *Sb-Kaf-CR1* | 33 | 21 (63.6%) | 1 (3.0%) |
| | | *Sb-Kaf-CR3* | | 22 (66.7%) | |

### Supplementary Table 10| *Sb-Lgs1* mutation and gene-dropout frequencies using two gRNAs in Macia

| Transformation system | Construct | sgRNA | # T0 plants analyzed | # of T0 events w/ mutation (freq.) | # T0 events w/ gene-dropout (freq.) |
|---|---|---|---|---|---|
| Integrated morphogenic gene-enabled | PHP87984 (*Axig1*$_{pro}$*:Wus2/Pltp*$_{pro}$*:Bbm*) | *Sb-Lgs1-CR1* | 42 | 35 (83.3%) | 22 (52.4%) |
| | | *Sb-Lgs1-CR3* | | 39 (92.9%) | |
| Non-integrated *Wus2*-enabled altruistic | (90%) PHP87980+(10%) PHP88158 (*3xENH:Pltp*$_{pro}$*:Wus2/NOS*$_{pro}$*:Crc*) | *Sb-Lgs1-CR1* | 161 | 65 (40.4%) | 47 (29.2%) |
| | | *Sb-Lgs1-CR3* | | 93 (57.8%) | |
